# Whole-mouse *in vivo* bioluminescence imaging applied to drug screening against *Leishmania infantum*: a reliable method to evaluate efficacy and optimize treatment regimens

**DOI:** 10.1101/326355

**Authors:** David M. Costa, Pedro Cecílio, Nuno Santarém, Anabela Cordeiro-da-Silva, Joana Tavares

## Abstract

Leishmaniasis is an important vector-borne neglected tropical disease caused by *Leishmania* parasites. Current anti-*Leishmania* chemotherapy is unsatisfactory, justifying the continued search for alternative treatment options. Herein, we propose the use of a minimally invasive bioluminescence-based murine model for preliminary *in vivo* screening of compounds against visceral infection by *Leishmania infantum*. We demonstrate that luciferase-expressing axenic amastigotes, unlike promastigotes, are highly infectious to BALB/c mice and generate a robust bioluminescent signal in the main target organs, such as the liver and spleen. Finally, we validate the use of this technique to evaluate *in vivo* treatment efficacy using reference drugs amphotericin B and miltefosine.

## MAIN TEXT

Leishmaniasis is a vector-borne parasitic disease caused by over 20 *Leishmania* species(1).It affects approximately 12 million people worldwide, with up to 1 million new cases every year (1). Visceral leishmaniasis (VL), the most severe form of the disease, is fatal if left untreated. VL is mainly associated with *Leishmania infantum* or *Leishmania donovani* infections, as these parasites are capable to disseminate to the host internal organs, particularly, the liver, spleen and bone marrow (1, 2). As there is still no vaccine available for humans, disease control relies mostly on chemotherapy and vector control. However, the limited and unsatisfactory chemotherapeutic options dictate the need of new drugs (3, 4). Indeed, every year up to 30 000 individuals suffering from VL die, some of them due to treatment failure (1, 5). Fortunately, neglected tropical diseases such as leishmaniasis have become a relevant part of the global health agenda, with a consequent increase in investment on control strategies (6). New leads against leishmaniasis are currently being optimized, while other compounds are already in pre-clinical and clinical stages (7, 8). Moreover, the recent development of *in vitro* high-throughput screening programs will undoubtedly feed the anti-*Leishmania* drug discovery pipeline with new compounds (8–10), whose efficacy remains to be addressed *in vivo*. Direct parasite observation remains the gold standard readout of anti-*Leishmania* drug *in vivo* efficacy (11). However, the traditional parasitological methods used to this end (microscopic observation of organ biopsies or limiting dilution assays) exhibit some limitations. Besides being labor-intensive and time-consuming, these methods only allow a static evaluation of infection since target organ collection entails euthanasia of the animal (8, 11). This is neither compatible with large scale-screening approaches nor ethically adequate, considering the requirement of a large number of animals (8). Thus, *in vivo* imaging techniques, namely those using bioluminescence-based models, have been developed to overcome such limitations. Nonetheless, these have either been mainly focused on cutaneous disease (8, 12) or require more than a month post-infection to warrant a readout (12–15). Here we show and validate a fast, non-invasive, bioluminescence-based mouse model of visceral infection by *L. infantum* suitable for an initial compound screening approach.

In a previous study, we demonstrated that luciferase-expressing *L. infantum* axenic amastigotes (16) injected intravenously generate a robust bioluminescent signal in mice (17). To assess if this signal could still be detected at later time points post-infection, thus allowing the assessment of treatment efficacy *in vivo*, we infected 6- to 7-week-old BALB/c mice with 10^8^ *L. infantum* axenic amastigotes by the intravenous (IV) route (Fig. 1A;C-D). Mice were then imaged 14 days post-infection (Fig. 1A) using an IVIS Lumina LT (PerkinElmer), 10 minutes after the subcutaneous administration of 2.4 mg of luciferin. The ventral fur was shaved to enable the maximization of detectable photons and the mice placed in dorsal position were angled to increase the detection of the signal coming from the spleen. Expectedly, the distribution of the bioluminescent signal indicates parasite establishment in the anatomical regions encompassing target organs such as the liver, spleen, lymph nodes and bone marrow (Fig. 1A). Interestingly, mice infected IV with the same inoculum of *L. infantum* promastigotes exhibited almost no detectable bioluminescent signal (Fig. 1B). Using the Living Image software, which can superimpose the bioluminescent signal of parasites and the grey-scale photograph of mice, elliptical regions (ROIs) were drawn to quantify bioluminescent signal in the anatomical regions of the liver, spleen, lymph nodes and bone marrow (the last two inferred from the signal of the left leg; Fig. 1A-B). At day 14 post-infection, the bioluminescent signal evaluated by the average radiance (photons/second/cm^2^/steradian) of the above ROIs was significantly higher in the animals infected with axenic amastigotes than in animals infected with promastigotes (Fig. 1C). Indeed, the signal in the spleen and leg of the animals infected with promastigotes was below the detection limit (Fig. 1C). To evaluate whether the differences in the bioluminescent signal detected in amastigote-and promastigote-infected mice were due to distinct infective capacities, parasite burden in the liver, spleen and bone marrow was evaluated using the gold standard limiting dilution assay (18). In fact, promastigote infection originated significantly lower parasite burdens in the liver, spleen and bone marrow when compared to axenic amastigote infection (Fig. 1D). This indicates that the difference in the signal intensity was most likely due to a reduced number of parasites in these organs. Since animals infected with axenic amastigotes yielded an early and sustained detectable bioluminescent signal in the main target organs, we used this model in a proof of concept experiment to validate it as a whole-animal imaging system to study the effectiveness of *in vivo* treatments against *L. infantum*. Consequently, infected animals were treated with miltefosine at 20 mg/kg/day (per os) or amphotericin B at 1 mg/kg/day (IV) for 4 days starting from day 15 post-infection (Fig. 2A). Imaging was performed right before treatment (day 15 post-infection), and one (day 19 post-infection) and three days (day 21 post-infection) after the last day of treatment (Fig. 2B). On day 21 post-infection mice were sacrificed and the liver, spleen and bone marrow were harvested for parasite burden assessment by limiting dilution. As anticipated, the short miltefosine treatment was sufficient to significantly decrease the bioluminescent signal in the ROIs defined for the liver and spleen (Fig. 2C). Amphotericin B was not as effective, although a statistically significant difference was still obtained in the spleen when compared to the untreated animals. Similar results were observed when the parasite burden was determined by the limiting dilution method (Fig. 2D). However, parasites were still detected in animals whose bioluminescent signal was bellow background levels (Fig. 2D). Therefore, despite lower sensitiveness, whole-animal bioluminescence imaging enabled the determination of the effectiveness of different treatments in reducing spleen and liver parasite burdens.

**FIG 1.**
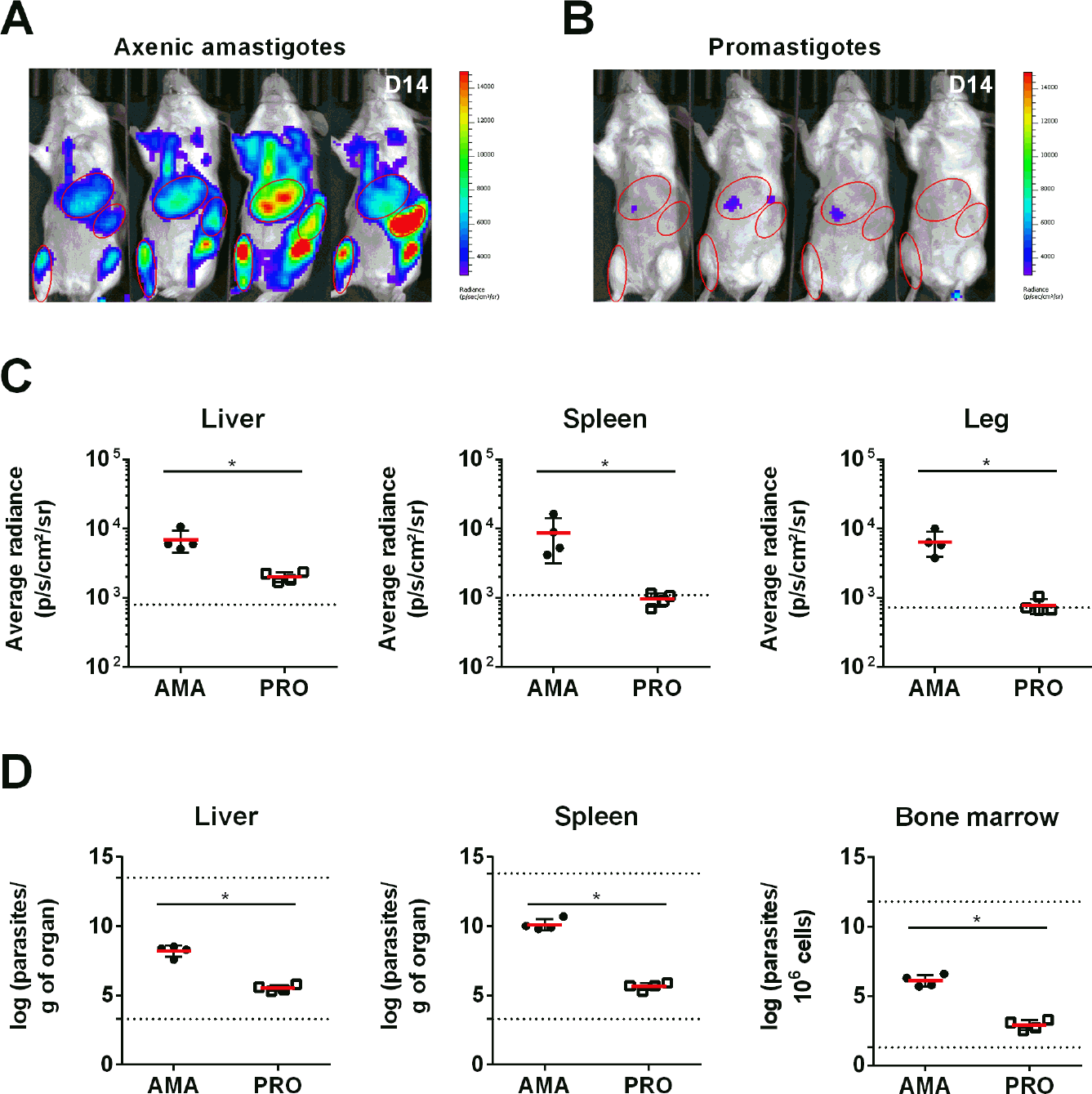
Infectivity of luciferase-expressing *L. infantum* axenic amastigotes and promastigotes via intravenous injection. (A, B) Images of BALB/c mice infected with either luciferase-expressing *L. infantum* axenic amastigotes (A) or promastigotes (B) resulting from the superimposition of the bioluminescence signal map and a grey-scale photograph of the mice. The regions of interest (ROIs) shown were used to quantify the bioluminescence signal originating from the liver, spleen, and right hind leg of the mouse. (C) Bioluminescence measurements expressed as average radiance (photons/s/cm^2^/steradian) corresponding to the previously defined liver (left), spleen (center), and right hind leg (right) ROIs. Means ± standard deviations are represented in bars. The dotted line represents the background signal calculated by applying the ROIs on images of uninfected animals. (D) Parasite burden in the liver, spleen, and femur 11 bone marrow determined by limiting dilution 14 days post-infection. Means ± standard deviations are represented in bars. The dotted lines represent the upper and lower detection limit of the technique for each organ. (C, D) AMA: animals infected with 10^8^ axenic amastigotes. PRO: animals infected with 10^8^ promastigotes. Statistical significance calculated by Mann Whitney test using Graphpad Prism 6.0 version: *p* <0.05 (*). Data representative of two independent experiments.

**FIG 2.**
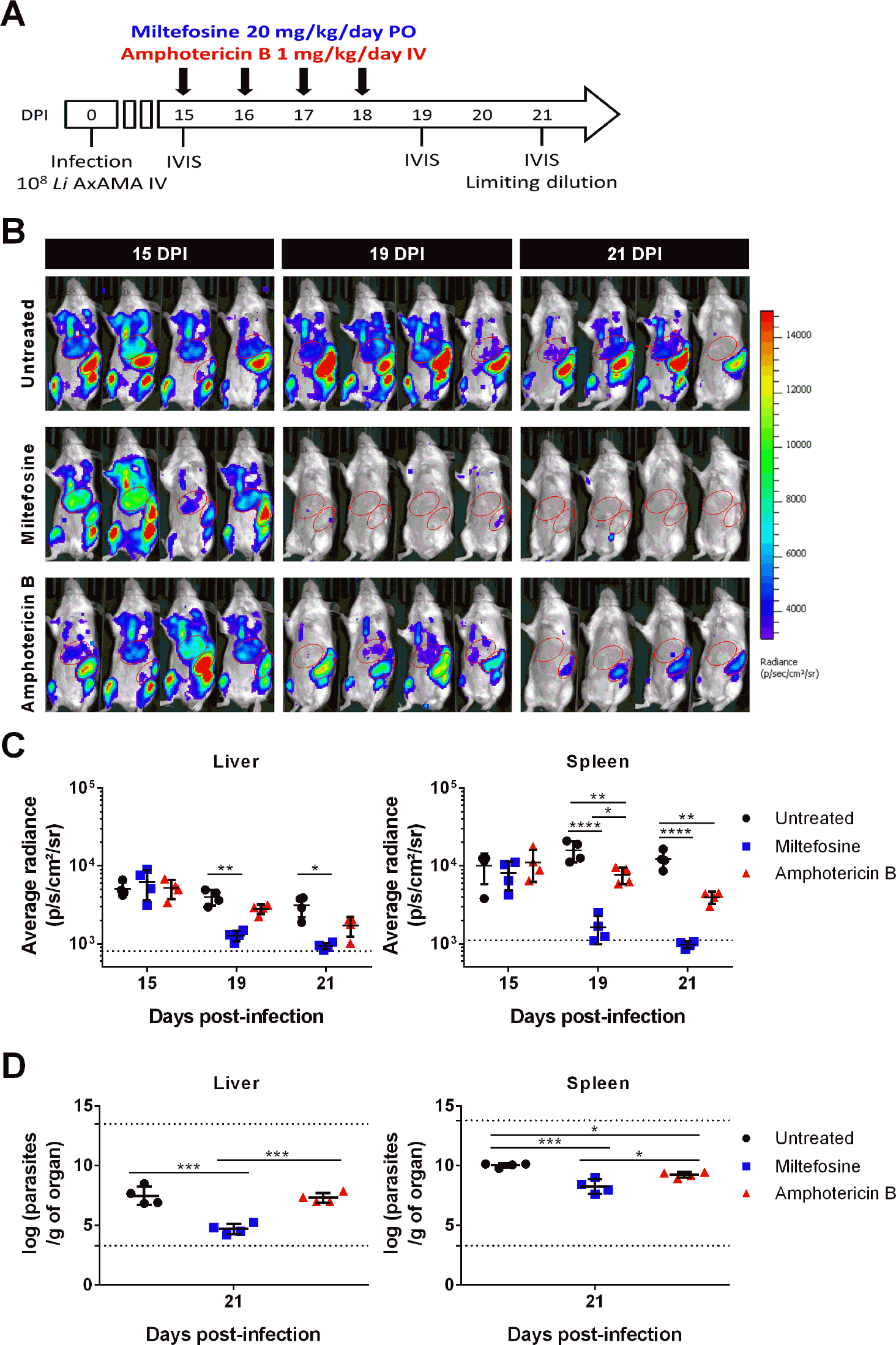
Treatment of *L. infantum* axenic amastigote-infected mice with reference drugs miltefosine and amphotericin B. (A) Schematic representation of the experimental design. BALB/c mice were infected with 10^8^ luciferase-expressing *L. infantum* axenic amastigotes (AMA) IV and 4-day treatments with either 20 mg/kg/day of miltefosine *per os* (PO) or 1 mg/kg/day of Amphotericin B IV were initiated 15 days post-infection 13 (DPI). All animals (n=4 per group) were imaged right before (day 15 post-infection), one day after (day 19 post-infection) and 3 days (day 21 post-infection) after the end of treatment using the IVIS Lumina LT system. At the last time point animals were sacrificed and parasite burden in the liver, spleen, and femur bone marrow were determined by limiting dilution. (B) Images of infected mice resulting from the superimposition of the bioluminescence signal map and a grey-scale photograph of the mice. The ROIs shown were used to quantify the bioluminescence signal originating from the liver and spleen anatomical regions. (C) Bioluminescence measurements expressed as average radiance (photons/s/cm^2^/steradian) corresponding to the previously defined liver (graph on the left) and spleen (graph on the right) ROIs. Means ± standard deviations are represented in bars. The dotted line represents the background signal calculated by applying the ROIs on images of uninfected animals. Statistical significance calculated by two-way ANOVA using Graphpad Prism 6.0 version: *p* <0.05 (*), *p<*0.005 (**), *p<*0.0001 (****). (D) Parasite burdens in the liver (graph on the left) and spleen (graph on the right) determined by limiting dilution 21 days post-infection. The dotted lines represent the upper and lower detection limit of the technique for each organ. Means ± standard deviations are represented in bars. Statistical significance calculated by ordinary one-way ANOVA using Graphpad Prism 6.0 version: *p<*0.05 (*), *p<*0.0005 (***). Data representative of two independent experiments.

We further evaluated the correlation between the two techniques used to determine parasite burdens in the liver and the spleen (Fig. 3). Average radiance values superior to the 99% confidence interval of the mean (Graphpad Prism 6.0 version) of the signal emitted by uninfected animals were plotted against the respective number of parasites per gram of liver (Fig. 3C) or spleen (Fig. 3D). Statistical significance, which translates into a positive correlation (Graphpad Prism 6.0 version), was found for both the liver and spleen determinations, evidencing the validity of our *in vivo* model.

**FIG 3.**
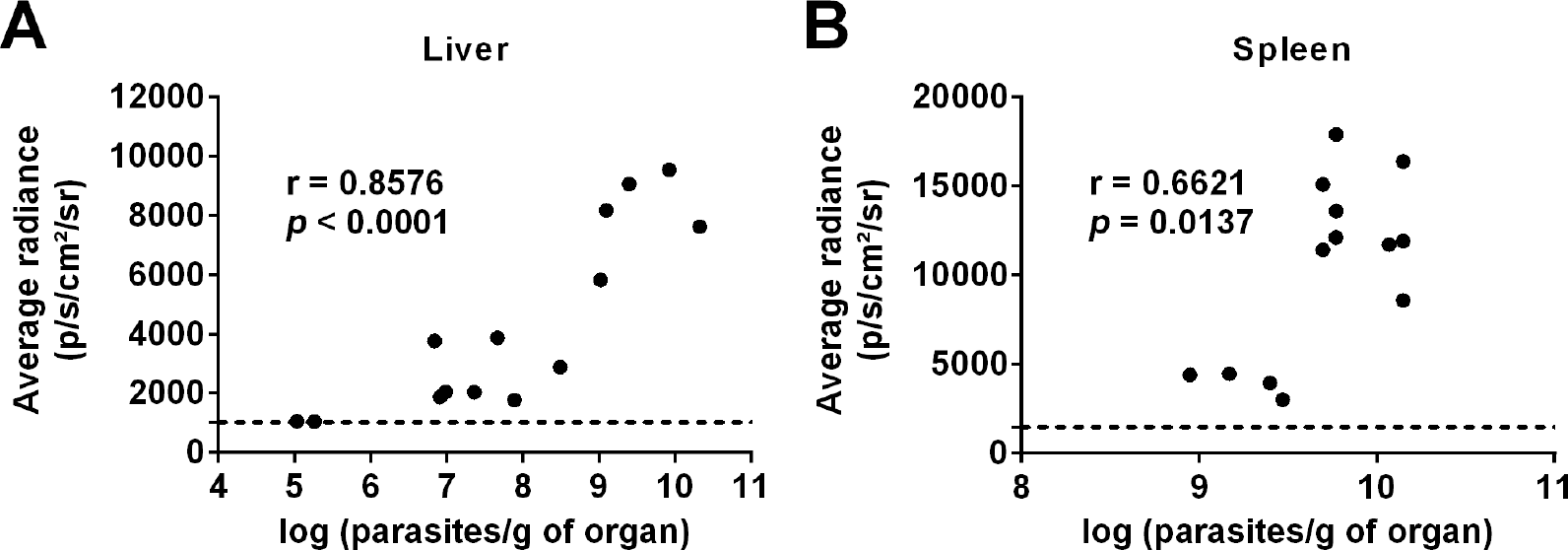
Relation between bioluminescence signal and parasite burdens measured by limiting dilution. Average radiance (photons/s/cm^2^/steradian) from liver (A) or spleen (B) ROIs plotted against the matching parasite burden in the corresponding organ. Pooled data of individual mice and from two independent experiments is shown. The dashed line represents the upper limit of the 99% confidence interval of the mean average radiance values obtained for each ROI when applied on images of uninfected BALB/c mice (n=6). Only animals displaying average radiance levels above the dashed line were considered for the calculation of the Pearson’s correlation coefficients using Graphpad Prism 6.0 version.

Using this model, spleen and liver parasite burdens remain stable during the first 4 weeks of infection (data not shown), leaving open the possibility of testing longer treatment regimens. Conversely, liver burdens predictably decrease to background bioluminescent levels at 8 weeks post-infection, suggesting the host could be controlling the infection due to granuloma formation in this organ [data not shown; (19)]. In contrast, spleen parasite burdens remain constant up to at least week 14, as evaluated by either bioluminescence imaging or limiting dilution assay (data not shown).

In conclusion, we propose the use of this rapid bioluminescence model for a preliminary *in vivo* screening of compounds against *L. infantum*. This minimally invasive method not only allows the accurate assessment of treatment efficacy, but also enables the adjustment of treatment regimens in an initial simple approach without the need to sacrifice large numbers of animals or to wait several days for a reliable readout. We expect this method to be a useful addition to the tools available to assist in the search for novel drugs to treat VL.

## ACKNOWLEDGMENTS

We thank Carla Oliveira from i3S for the support with the statistical analysis.

## FUNDING

This work was supported by funds from the Fundação para a Ciência e Tecnologia (FCT)/Ministério da Educação e Ciência (MEC) co-funded by the European Regional Development Fund (FEDER) under the Partnership agreement PT2020, through the Research Unit No.4293. This work also received funds from project Norte-01-0145- FEDER-000012 - Structured program on bioengineered therapies for infectious diseases and tissue regeneration, supported by Norte Portugal Regional Operational Programme (NORTE 2020), under the PORTUGAL 2020 Partnership Agreement, through the FEDER. JT is an Investigator FCT funded by National funds through FCT and co-funded through European Social Fund within the Human Potential Operating Programme. DC and PC are funded by FCT through individual fellowships (SFRH/BD/123734/2016 and SFRH/BD/121252/2016, respectively).

## REFERENCES

1. WHO. 2017 Leishmaniasis, Fact sheet No 375 http://www.who.int/mediacentre/factsheets/fs375/en/. Accessed January 10, 2018

2. Cecilio P, Perez–Cabezas B, Sanatarem N, Maciel J, Rodrigues V, Cordeiro da Silva A. 2014. Deception and manipulation: the arms of leishmania, a successful parasite. Front Immunol 5:480.

3. Loureiro I, Faria J, Sanatarem N, Smith TK, Tavares J, Cordeiro-da-Silva A. 2017. Potential drug targets in the pentose phosphate pathway of trypanosomatids. Curr Med Chem dio:10.2174/0929867325666171206094752.

4. Ponte-Sucre A, Gamarro F, Dujardin JC, Barrett MP, Lopez-Velez R, Garcia-Hernandez R, Pountain AW, Mwenechanya R, Papadopoulou B.2017. Drug resistance and treatment failure in leishmaniasis: A 21st century challenge. PLoS Negl Trop Dis 11:e0006052.

5. Moore EM, Lockwood DN. 2010. Treatment of visceral leishmaniasis. J Glob Infect Dis 2:151–158.

6. Molyneux DH, Savioli L, Engels D. 2017. Neglected tropical diseases: progress towards addressing the chronic pandemic. Lancet 389:312–325.

7. Field MC, Horn D, Fairlamb AH, Ferguson MA, Gray DW, Read KD, De Rycker M, Torrie LS, Wyatt PG, Wyllie S, Gilbert IH. 2017. Anti-trypanosomatid drug discovery: an ongoing challenge and a continuing need. Nat Rev Microbiol 15:217–231.

8. Zulfiqar B, Shelper TB, Avery VM. 2017. Leishmaniasis drug discovery: recent progress and challenges in assay development. Drug Discov Today 22:1516–1531.

9. Annang F, Perez-Moreno G, Garcia-Hernandez R, Cordon-Orbas C, Martin J, Tormo JR, Rodriguez L, de Pedro N, Gomez-Perez V, Valente M, Reyes F, Genilloud O, Vicente F, Castanys S, Ruiz-Perez LM, Navvaro M, Gamarro F, Gonzalez-Pacanowska D. 2015. High-throughput screening platform for natural product-based drug discovery against 3 neglected tropical diseases: human African trypanosomiasis, leishmaniasis, and Chagas disease. J Biomol Screen 20:82–91.

10. Pena I, Pilar Manzano M, Cantiazani J, Kessler A, Alonso-Padilla J, Bardera AI, Alvarez E, Colmenarejo G, Cotillo I, Roquero I, de Dios-Anton F, Barroso V, Rodriguez A, Gray DW, Navvaro M, Kumar V, Sherstnev A, Drewry DH, Brown JR, Fiandor JM, Julio Martin J. 2015. New compound sets identified from high throughput phenotypic screening against three kinetoplastid parasites: an open resource. Sci Rep 5:8771.

11. Gupta S, Nishi. 2011. Visceral leishmaniasis: experimental models for drug discovery. Indian J Med Res 133:27–39.

12. Avci P, Karimi M, Sadasivam M, Antunes-Melo WC, Carrasco E, Hamblin MR. 2017. In-vivo monitoring of infectious diseases in living animals using bioluminescence imaging. Virulence doi:10.1080/21505594.2017.1371897:1–35.

13. Melo GD, Goyard S, Lecoeur H, Rouault E, Pescher P, Fiette L, Boissonnas A, Minoprio P, Lang T.2017. New insights into experimental visceral leishmaniasis: Real-time in vivo imaging of Leishmania donovani virulence. PLoS Negl Trop Dis 11:e0005924.

14. Michel G, Ferrua B, Lang T, Maddugoda MP, Munro P, Pomares C, Lemichez E, Marty P.2011. Luciferase-expressing Leishmania infantum allows the monitoring of amastigote population size, in vivo, ex vivo and in vitro. PLoS Negl Trop Dis 5:e1323.

15. Reimao JQ, Oliveira JC, Trinconi CT, Cotrim PC, Coelho AC, Uliana SR. 2015. Generation of luciferase-expressing Leishmania infantum chagasi and assessment of miltefosine efficacy in infected hamsters through bioimaging. PLoS Negl Trop Dis 9:e0003556.

16. Sereno D, Roy G, Lemesre JL, Papadopoulou B, Ouellette M. 2001. DNA transformation of Leishmania infantum axenic amastigotes and their use in drug screening. Antimicrob Agents Chemother 45:1168–1173.

17. Tavares J, Costa DM, Teixeira AR, Cordeiro-da-Silva A, Amino R. 2017. In vivo imaging of pathogen homing to the host tissues. Methods 127:37–44.

18. Buffet PA, Sulahian A, Garin YJ, Nassar N, Derouin F. 1995. Culture microtitration: a sensitive method for quantifying Leishmania infantum in tissues of infected mice. Antimicrob Agents Chemother 39:2167–2168.

19. Rodrigues V, Cordeiro-da-Silva A, Laforge M, Silvestre R, Estaquier J. 2016. Regulation of immunity during visceral Leishmania infection. Parasit Vectors 9:118.

